# 3D imaging of the pregnant uterus reveals an extensively invasive mouse placenta and early CXCL12-CXCR4 requirement

**DOI:** 10.64898/2026.04.06.716785

**Authors:** James B. Zwierzynski, Mira N. Moufarrej, Kristy Red-Horse

## Abstract

Successful pregnancy requires exquisite balance: the placenta must invade just enough to access maternal blood but not so deep it remains attached at birth. Disrupting this balance causes life-threatening pregnancy complications, for which treatments remain limited. Animal models are desperately needed to discover mechanisms underlying balanced uteroplacental development and how pregnancy complications arise, but this is hampered by the view that mouse placentation lacks human characteristics such as extensive trophoblast invasion and targeting of uterine spiral arteries. Here, we utilize 3D imaging, mouse genetics, and pharmacological perturbations to demonstrate that: (1) The mouse placenta invades more extensively than previously recognized with most spiral arteries heavily enveloped by fetal trophoblasts, (2) This process is disrupted without CXCL12-CXCR4 signaling specifically during early pregnancy, and (3) Disrupting early uteroplacental development ultimately results in excessively deep trophoblast invasion, closely mimicking the pregnancy complication placenta accreta. Mechanistically, uterine epithelium, stroma, and arteries activate CXCR4 signaling in early pregnancy, and inhibition causes decidualization failure, followed by dissolution of spiral artery development. Trophoblasts consequently migrate deep into uterine muscle and its arteries, reproducing hallmarks of human accreta. Thus, with 3D imaging, the mouse more effectively models human uteroplacental development and defines an early etiological window for intervention.

## Main

During pregnancy, complex interactions between maternal cells and placental trophoblasts transform the uterine wall into an environment that supports the developing fetus, while at the same time maintaining maternal health.^1–3^ Maternal spiral arteries must grow and remodel within the uterine endometrial layer, and the resident stromal cells must undergo a cellular and molecular transformation called decidualization (e.g., expansion and expression of key genes such as *Prl8a2, Cdh3, Igfbp1*), which provides a niche for trophoblasts to invade and tap into a maternal arterial blood supply.^4–8^ Here, trophoblasts must migrate deeply enough to form a firm decidual attachment and access spiral arteries, but they must avoid invading the deeper myometrium so the placenta can safely separate at birth. Pregnancy complications arise when this process is not properly balanced; shallow invasion is hypothesized to cause preeclampsia and fetal growth restriction^9–11^ while excessive invasion is associated with placenta accreta.^12^ Although these complications are detected and diagnosed in mid-to-late pregnancy, the prevailing model states that they arise from aberrant trophoblast invasion and artery transformation in early pregnancy.^10,13–17^ Despite this hypothesis, a lack of animal models hinders our fundamental understanding of normal uteroplacental development and the precise sequence of events underlying pathological pregnancies.

Placenta accreta is a life-threatening condition where the placenta develops abnormal adhesion so that it does not fully detach after delivery, leading to maternal hemorrhage or severe infection, with hysterectomy as the standard of care.^12^ It complicates nearly 1 in 250 births in the United States, a 470% increase over the past 50 years, driven largely by rising cesarean sections rates.^18,19^ Based on ultrasound and histological observations, the primary cause is hypothesized to be a thin, absent, or defective decidua, thereby setting the stage for trophoblast invasion into, and even through the myometrium, to seek an alternative blood supply.^17,20–23^ Through surgeries simulating cesarean section sequelae or genetic knockouts (*Adm*, *Marveld1*, *Gab3*) in mice, researchers have modeled some placenta accreta-like features.^24–29^ However, capturing the full developmental complexity of human placenta accreta, particularly whether specific molecular disruptions during early pregnancy cause pathology months later, remains a formidable challenge. Without such evidence, and without robust animal models, the sequence of events linking early defects with later pathology remains theoretical, and opportunities for early intervention remain out of reach.

The mouse represents one of the most genetically tractable mammalian models and shares some key features with human pregnancy. Both species induce decidualization and develop decidual spiral arteries that supply a hemochorial placenta with blood, although differences exist in timing relative to implantation.^4^ Despite these similarities, murine pregnancy has long been considered an inadequate model of human pregnancy owing to perceived shallow trophoblast invasion and limited spiral artery targeting.

Literature suggests a lack of bulk trophoblast invasion, positing that cells primarily localize perivascularly at the midline of the placenta with little interstitial and peripheral invasion.^30–32^ Consequently, researchers are frequently hesitant to use the mouse to perform *in vivo* studies of human disease-relevant characteristics of the maternal fetal interface.

However, these perceptions have been shaped predominantly by two-dimensional (2D) histological analyses of tissue sections, which do not always comprehensively capture three-dimensional (3D) vascular and cellular relationships.^33–36^ In a variety of organs and model systems, including zebrafish, mice, and human tissue, 3D imaging has been key to illuminate brain vascular and meningeal anatomy,^33,35,37,38^ artery development through an initial remodeling zone,^39^ potential regenerative targets,^40–45^ and artery patterning through secreted CXCL12.^46^ We hypothesized that applications of 3D imaging to understand uteroplacental development and disease could be similarly fruitful.

Using whole-organ 3D imaging across gestation, we demonstrate that the mouse placenta invades extensively—thousands of trophoblasts migrate through the entire decidual thickness. Comprehensive quantification of the maternal-fetal interface, and temporal lineage tracing, revealed that spiral arteries differentiate from local capillaries and develop proximally to the placenta shortly after implantation. Subsequently, trophoblasts extensively invade, both interstitially and perivascularly, where they envelop three-quarters of the maternal spiral arteries, connecting them to the placenta.

This 3D analysis platform was used to interrogate a secreted ligand-receptor axis, CXCR12/CXCR4,^47–50^ that is important for the development of mouse fetal umbilical arteries and has been predicted in spatial omics studies to function at the human maternal-fetal interface.^51–53^ We found mouse *Cxcl12* to be expressed by trophoblasts during a short window post-implantation at a time when adjacent uterine epithelium, stroma, and arteries all activate CXCR4 signaling. Fetal deletion of *Cxcl12* or pharmacologic CXCR4 inhibition, specifically during early pregnancy, results in decidualization failure, dissolution of nascent arteries, and deep trophoblast invasion into the myometrium and its arteries. By late pregnancy, these defects result in hallmarks of placenta accreta including uterine adhesions and hemorrhage.

In total, we demonstrate that comprehensive analysis of the maternal-fetal interface in 3D (1) reveals disease-relevant human features in the mouse, (2) enables us to define the role of CXCL12-CXCR4 signaling in early pregnancy establishment, and (3) provides a new model of placenta accreta.

## Results

### 3D imaging reveals a highly invasive murine placenta

To obtain a more holistic view of trophoblast invasion and their interactions with maternal arteries during mouse pregnancy (Fig 1a), we harvested a time course of non-pregnant and pregnant uteri and performed whole-organ immunolabeling for trophoblasts (CK8) and arteries (SMA/JAG1) followed by clearing and light sheet microscopy. Concomitantly, conventional analyses were performed where these same antibodies were used to label tissue sections from the midline of the placenta.^30,31^ Note that CK8 also stains uterine luminal epithelium and the mesometrial covering of the uterus, but their distinct morphology and spatial localization allow clear differentiation from trophoblasts in our images.

**Figure 1:**
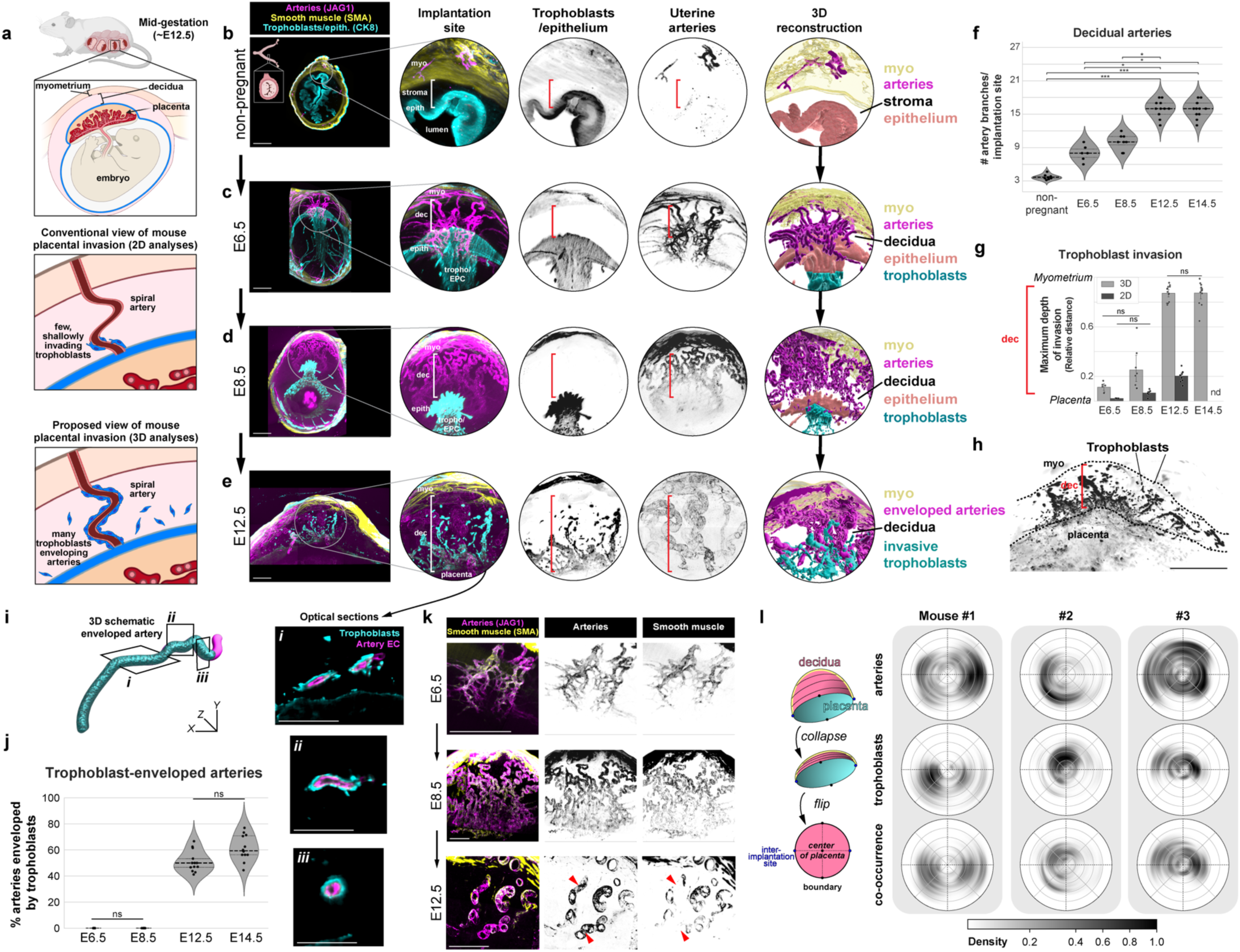
3D imaging reveals a highly invasive murine placenta. **a**, Schematic of mouse placental anatomy and proposed overviews of mouse placental invasion. **b-e,** Representative cross-sectional 200µm maximum intensity projections and 3D reconstructions from whole-organ light sheet imaging of the non-pregnant mouse uterus (**b**), and implantation sites at E6.5 (**c**), E8.5 (**d**), and E12.5 (**e**). **f**, Quantification of artery branches entering the endometrium from the myometrium, across whole implantation sites, by whole-organ light sheet imaging. Mice were quantified at non-pregnancy (n = 9 mice), E6.5 (n = 6 implantation sites, n = 3 mice), E8.5 (n = 8 implantation sites, n = 4 mice), E12.5 (n = 13 implantation sites, n = 4 mice), and E14.5 (n = 12 implantation sites, n = 3 mice). **g**, The maximum relative distance of trophoblast invasion from the placenta to the myometrium across gestation. Mice were quantified in 3D at: E6.5 (n = 6 implantation sites, n = 3 mice), E8.5 (n = 8 implantation sites, n = 4 mice), E12.5 (n = 13 implantation sites, n = 4 mice), and E14.5 (n = 12 implantation sites, n = 3 mice). Mice were quantified in 2D at: E6.5 (n = 4 implantation sites, n = 2 mice), E8.5 (n = 7 implantation sites, n = 2 mice), E12.5 (n = 10 implantation sites, n = 3 mice); nd, no data collected. All comparisons not denoted as ns are significant *P* < 0.01. **h**, Maximum projection of CK8 in the decidua of an E12.5 implantation site. **i**, 3D schematic of a spiral artery enveloped by trophoblasts and 5µm optical sections from light sheet microscopy (i-iii). **j,** Quantification of the percentage of trophoblast-enveloped arteries. Mice were quantified at E6.5 (n = 6 implantation sites, n = 3 mice), E8.5 (n = 8 implantation sites, n = 4 mice), E12.5 (n = 13 implantation sites, n = 4 mice), and E14.5 (n = 12 implantation sites, n = 3 mice). All comparisons not denoted on graph are *P* < 0.05. **k**, Representative 200µm optical sections of arterial smooth muscle localization at E6.5, E8.5, and E12.5 on spiral arteries. **l**, Schematic for the construction of 3D probability densities (left). Plots (right) for 3D localization of arteries and trophoblasts within the decidua and myometrium (n = 3 implantation sites, n = 3 mice). \**P* <0.05, \*\**P* <0.01, \*\*\**P* <0.001. *P* values from Dunn Post-Hoc test with Bonferroni Correction (**f**, **j**), Independent t-test (**g**, within gestational age), or One-way ANOVA with Tukey Post-Hoc Test (**g,** across imaging modality). Scale bars, 500µm. myo, myometrium; epith, epithelium; dec, decidua; tropho/EPC, trophoblasts/ectoplacental cone.

We found that before pregnancy, uterine arteries primarily reside within the outer muscular layer of the uterus called the myometrium, with artery branches rarely extending into the endometrial stroma (Fig 1b, f), and CK8 labeling the epithelium (Fig 1b). In contrast, 2 days after embryo implantation at embryonic day (E) 6.5, an average of 8 individual arteries had formed in the decidualized stroma (Fig 1c, f, Supplementary Movie 1). These new arteries loosely spiraled directly toward the localized site of proliferating trophoblasts, which at this stage are called the ectoplacental cone, in an arrangement reminiscent of the nascent artery remodeling zones observed during embryonic artery development.^39^ Trophoblasts had yet to extensively invade at this time point with a few individual trophoblasts found approximately 10% of the way through the decidua (Fig 1g, Extended Data 1a, left, arrowheads). As pregnancy progresses over the next two days, at E8.5, the artery network quickly matured forming the expected coiled pattern of spiral arteries with an average of 10 branching from the myometrium (Fig 1d, f, Supplementary Movie 2). A few trophoblasts had also advanced to 25% of the distance migrated through the decidua (Fig 1g, Extended Data Fig 1a, right, arrowheads). Thus, following implantation, numerous spiral arteries develop in the mouse decidua descending from the myometrium toward the ectoplacental cone, occurring prior to most trophoblast invasion.

By mid-pregnancy, at E12.5, we observed a typical mature placenta morphology with an average of 16 spiral arteries per implantation site (Fig 1e, f, Supplementary Movie 3). There was also a dramatic increase in trophoblast invasion at this time point; trophoblasts had now invaded nearly 90% of the way through the decidua to the myometrium (Fig 1e, g). When observing a maximum projection of the decidua, thousands of trophoblasts were visible up to the myometrial border (Fig 1h).

Remarkably, approximately 50-75% of spiral arteries were enveloped by invading trophoblasts through most of their length (Fig 1i, j). All enveloped arteries connected to the placenta (Extended Data Fig 1b), and many exhibited patchy loss of smooth muscle (Fig 1k, arrowheads), suggesting these substantial artery-trophoblast interactions may play a role in artery transformation and be critical in shunting blood to the developing embryo. These general anatomical formations were also observed in tissue sections at the midline of the placenta (Extended Data Fig 1c-f), but the extent of trophoblast invasion is highly underestimated with this method. Quantification revealed an average 4–5-fold discrepancy (Fig 1g). Thus, 3D microscopy highlights previously underappreciated features of the murine maternal-fetal interface and enables quantification of complex vascular and cellular relationships.

We next sought to further understand the discrepancy between 2D and 3D measurements. To do so, we extracted the 3D positions of arteries and trophoblasts and constructed their probability distributions within the half-dome-like region of the decidua and myometrium, excluding the mesometrium where CK8 signal from trophoblasts and epithelium was indistinguishable (see Methods) (Extended Data Fig 1g, Supplementary Movie 4) We found that arteries were generally centralized within the decidua, but trophoblast invasion displayed random bias, often skewed in one direction (Fig 1l).

Consistent with these patterns, artery and trophoblast co-occurrence is constrained by the presence of trophoblasts (Fig 1l) and clarified our observation that trophoblast-enveloped arteries were not consistently centrally located (Supplemental Movie 3).

Thus, holistically studying trophoblast-artery interactions within the decidua necessitates 3D whole-organ imaging since analyses that rely on centrally sampled 2D tissue sections underestimates the degree and artery-bias of trophoblast invasion owing to its unpredictability in 2D space. These data provide a comprehensive 3D map of murine uteroplacental development, demonstrating that the mouse maternal-fetal interface, like human, displays deep trophoblast invasion and their strong tropism for spiral arteries (Fig 1a).

### Spiral arteries develop in early pregnancy when CXCL12-CXCR4 signaling is active

Our 3D map illuminated the structural complexity of spiral artery growth, yet the developmental trajectory to create this network remained unclear. To understand how these features emerge, we asked: How do spiral arteries grow to supply blood to the placenta? What are their cellular origins? Is their differentiation limited to a certain developmental timepoint? To investigate these questions, we first tested whether spiral arteries develop via a commonly implemented mechanism of arterial formation – the differentiation of capillary and venous endothelial cells (ECs) into artery ECs.^41, 54–59^ This can be tested using lineage tracing with an *ApjCreER* mouse line that, when crossed with the Cre reporter, *Rosa26^tdTomato^,* labels capillary and venous ECs but not arteries (Fig 2a). To ensure cellular resolution, we utilized tissue sections and confocal imaging to quantify this lineage tracing. Injecting non-pregnant *ApjCreER; Rosa26^tdTomato^* mice with tamoxifen one day before harvesting the uterus to assess initial cell labeling confirmed almost exclusive labeling of capillary and venous endothelium with only rare artery ECs being labeled (Extended Data Fig 2a). Extending the tracing period from pre-pregnancy through E12.5 revealed that all spiral arteries indeed differentiate almost exclusively from capillary and venous progenitors (Fig 2b, Extended Data Fig 2b).

**Figure 2:**
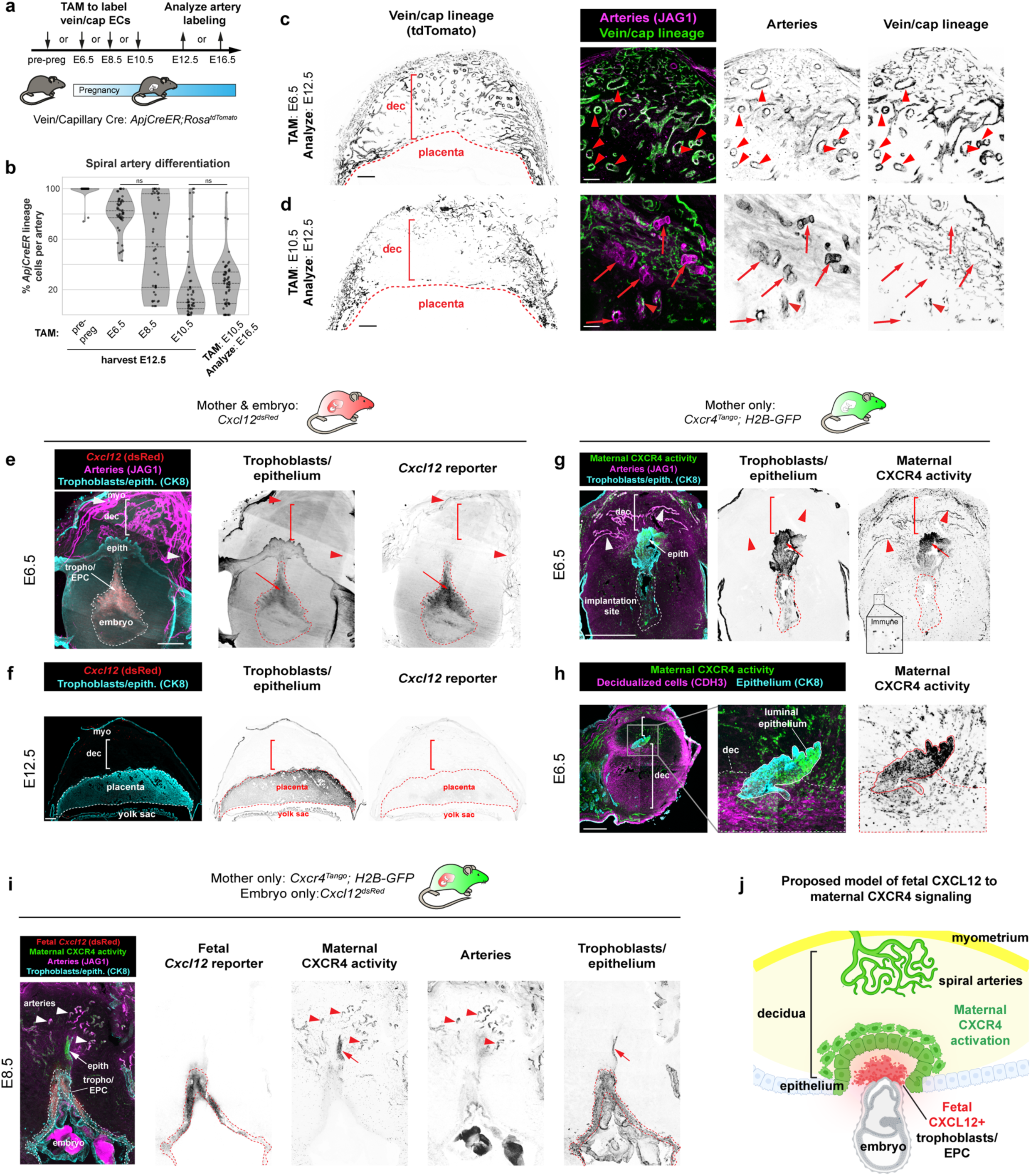
Spiral arteries develop early in pregnancy when CXCL12-CXCR4 signaling is active. **a**, Experimental strategy for (**b-d**). **b**, Quantification of *ApjCreER* lineage labeling of arteries. Mice were quantified at E12.5 from inductions at pre-pregnancy (n = 6 implantation sites, n = 3 mice), E6.5 (n = 6 implantation sites, n = 3 mice), E8.5 (n = 5 implantation sites, n = 3 mice), E10.5 (n = 5 implantation sites, n = 3 mice), and quantified at E16.5 from induction at E10.5 (n = 6 implantation sites, n = 3 mice). All *P* values not denoted as n.s. < 0.001. **c**, *ApjCreER* lineage labeling of spiral arteries at E12.5 with E6.5 induction. Arrowheads denote lineage labeling of arteries. **d**, *ApjCreER* lineage labeling of spiral arteries at E12.5 with E10.5 induction. Arrowhead denotes cluster of lineage labeled cells, arrows indicate spiral arteries lacking lineage labeling. **e**, Representative cross-sectional 60µm optical sections from whole-organ light sheet imaging of *Cxcl12^dsRed^* at E6.5. Arrowheads denote maternal arteriole *Cxcl12* expression and arrow denotes trophoblastic *Cxcl12* expression. **f**, Representative immunofluorescence of *Cxcl12^dsRed^* at E12.5. **g**, Immunofluorescence of maternal CXCR4 activation at E6.5 in maternal arteries (arrowheads) and epithelium, and **h**, in decidualized stroma. **i**, Immunofluorescence of fetal *Cxcl12* and maternal CXCR4 activation at E8.5. **j**, Schematic for a potential role of CXCL12 and CXCR4 in early pregnancy. *P* values from Dunn Post-Hoc test with Bonferroni Correction. Scale bars: 100µm, **c**,**d**, right three panels; 500µm, all others. dec stroma, decidualized stroma; myo, myometrium; tropho, trophoblasts; epith, epithelium; EPC, ectoplacental cone.

We then performed windows of lineage tracing to identify the developmental timepoint during which spiral arteries differentiated from their vein/capillary progenitors. Labeling at E6.5 resulted in the majority of E12.5 spiral artery ECs traced (Fig 2b, c, arrowheads), while E10.5 labeling resulted in much less tracing, and the traced arteries mostly contained only single ECs or small clusters (Fig 2b, d, arrowheads, untraced arteries; arrow, traced ECs). When E10.5 labeling was extended to E16.5, the tracing percentage was similar to the E10.5-12.5 window (Fig 2b, Extended Fig 2b). These data indicated that most spiral artery differentiation occurs in the early phase of pregnancy before E10.5, which is consistent with our finding that artery branching into the decidua plateaus by E12.5 (Fig 1f).

To provide further support, we also conducted an opposing lineage tracing experiment, where artery ECs were labeled with *Cx40CreER; Rosa26^tdTomato^*mice at E10.5 and analyzed using the same methods to test whether labeled spiral artery cells were replaced at E12.5. Most spiral arteries remained labeled at E12.5, indicating that they were not substantially replaced (Extended Data Fig 2c, arrowheads). Thus, our complementary lineage tracing experiments demonstrate that spiral artery differentiation occurs primarily before E10.5 through differentiation of the vein/capillary EC population.

Because of its known role in artery development,^40,41,46–51,60–65^ we investigated whether CXCL12-CXCR4 signaling was active during spiral artery development in mice. We first mapped the expression of *Cxcl12* and activation of CXCR4 throughout gestation using the *Cxcl12^dsRed^* and *Cxcr4^Tango^* reporters.^50,66^ In early pregnancy, at E6.5 and E8.5, *Cxcl12* expression was detected in trophoblasts of the ectoplacental cone and maternal myometrial arteries, but not in spiral arteries in the decidua (Fig 2e, arrow, trophoblasts; arrowheads, myometrial arteries; Extended Data 3a). However, trophoblast expression was downregulated by E12.5 (Fig 2f). Therefore, fetal trophoblasts of the ectoplacental cone express *Cxcl12* at the time points coinciding with the bulk of spiral artery growth, but expression is turned off after this early time window.

To gain insight to which cells respond to CXCL12, we used a CXCR4 activity reporter *Cxcr4^Tango^; pTRE-H2B-GFP,* which expresses H2B-GFP in cells containing an activated CXCR4.^50,67,68^ At E6.5, ECs of maternal spiral arteries differentiating proximally to the ectoplacental cone and immune cells throughout the implantation site exhibited CXCR4 activation (Fig 2g, arrowheads, spiral arteries, Extended Data Fig 3b, arrowheads). Unexpectedly, CXCR4 was also strongly activated on maternal luminal epithelium (Fig 2g, h) and nearby CDH3+ decidual stromal cells (Fig 2h), but only in the region directly above the *Cxcl12*-expressing trophoblasts of the ectoplacental cone (Fig 2i). CXCR4 activity was never observed on invasive trophoblasts (Extended Data Fig 3c).

Together, our analyses identify the progenitors of uterine spiral arteries in mice and demonstrate that their differentiation primarily occurs during the period of transient trophoblastic *Cxcl12* expression in early pregnancy. Activation of CXCR4 in the maternal epithelium and decidua surrounding the ectoplacental cone at this same timepoint suggests that trophoblast CXCL12 may play a role in decidualization, while concurrent activity in spiral arteries may be a chemotactic signal to guide arterial growth (Fig 2j).

### Early fetal CXCL12 is required for uteroplacental development

Next, we sought to test for the role of trophoblast *Cxcl12* in spiral artery development and trophoblast invasion using both genetic and pharmacological tools (Fig 3a).

**Figure 3:**
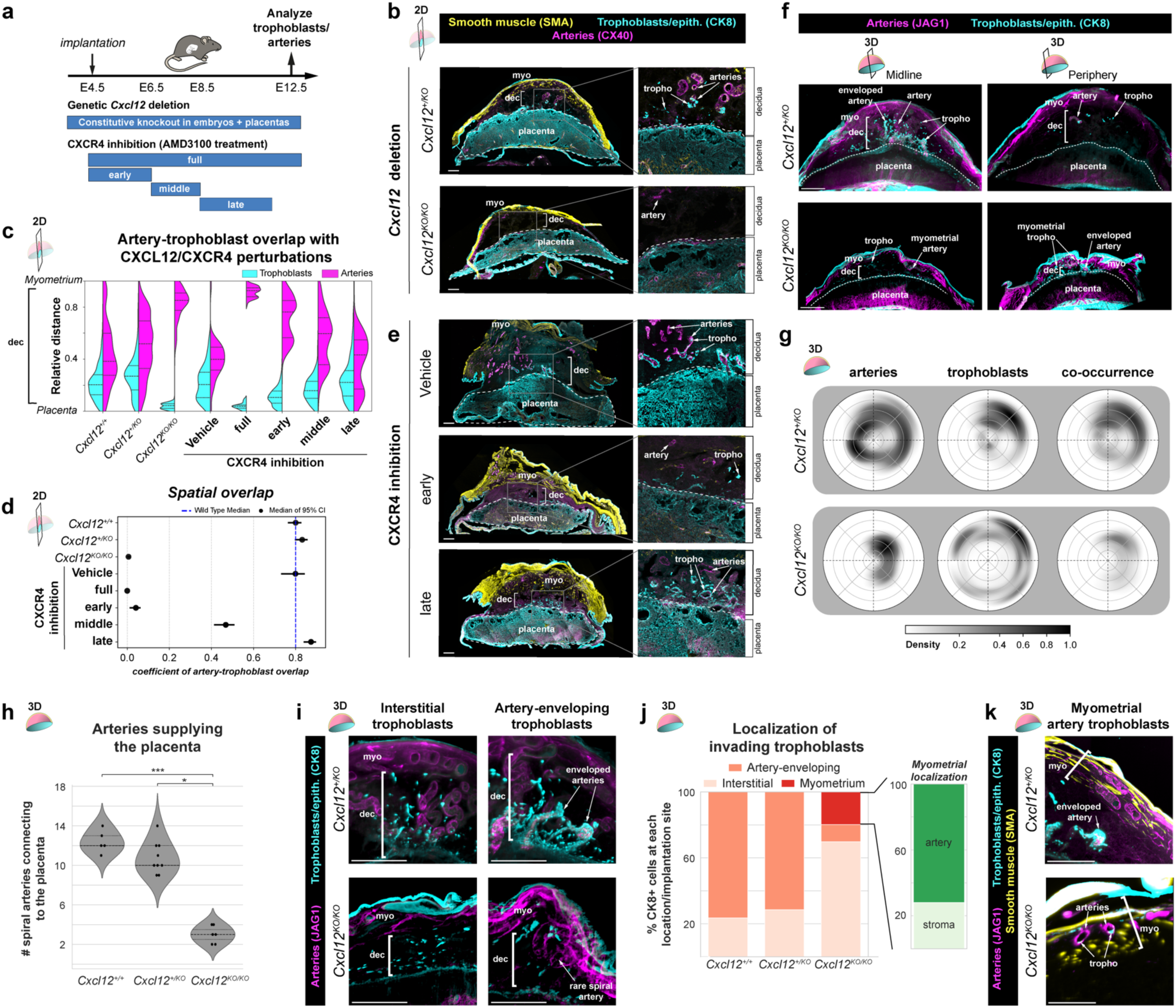
Early fetal CXCL12 is required for uteroplacental development. **a**, Genetic and pharmacological approaches for implantation site phenotyping. **b**, Immunofluorescence of E12.5 implantation sites from *Cxcl12^+/KO^* intercrosses to examine artery-trophoblast interactions in the uterine wall. **c**, Trophoblast and artery positions relative to the placenta-myometrium distance. **d**, Odds ratios with 95% confidence intervals for the interaction of artery and trophoblast interactions in the uterine wall. **e**, Immunofluorescence of E12.5 implantation sites treated with CXCR4 inhibitor to examine artery-trophoblast interactions in the uterine wall. **f**, Representative cross-sectional 200µm optical sections from whole-organ light sheet imaging of E12.5 *Cxcl12^+/KO^* and *Cxcl12^KO/KO^* mouse implantation sites in the midline and periphery of the implantation sites. **g**, Plots for 3D localization of arteries and trophoblasts within the decidua and myometrium. **h**, Quantification of number of maternal arteries entering the placenta from whole-organ light sheet imaging. **i**, 100µm optical sections of interstitial and spiral-artery associated trophoblasts in from *Cxcl12^+/KO^* intercrosses at E12.5. **j**, Quantification of trophoblast localization from whole-organ light sheet imaging from *Cxcl12^+/KO^* intercrosses at E12.5. **k**, 100µm optical sections of myometrial artery trophoblasts in *Cxcl12^KO/KO^* implantation sites, arrowheads denote trophoblasts in myometrial artery. Mice analyzed in 2D at E12.5, *Cxcl12*^+/+^ (n = 9 implantation sites, n = 3 mice), *Cxcl12^+/KO^* (n = 11 implantation sites, n = 3 mice), *Cxcl12^KO/KO^* (n = 18 implantation sites, n = 5 mice), Vehicle (n = 6 implantation sites, n = 3 mice), AMD E4.5-11.5 (n = 10 implantation sites, n = 3 mice), AMD E4.5-6.5 (n = 14 implantation sites, n = 3 mice) AMD E6.5-8.5 (n = 13 implantation sites, n = 3 mice), AMD E8.5-11.5 (n = 11 implantation sites, n = 3 mice). Mice analyzed in 3D at E12.5, *Cxcl12^+/+^* (n = 5 implantation sites, n = 3 mice, *Cxcl12^+/KO^* (n = 9 implantation sites, n = 3 mice), *Cxcl12^KO/KO^* (n = 7 implantation sites, n = 3 mice). \**P* <0.05, \*\**P* <0.01, \*\*\**P* <0.001. *P* values from Dunn Post-Hoc test with Bonferroni Correction. Scale bars, 500µm. myo, myometrium; tropho, trophoblast; dec, decidua.

Replacing an exon on *Cxcl12* with a dsRed cassette generates a knockout allele, hereafter referred to as *Cxcl12^KO^*. Intercrosses between *Cxcl12^+/KO^* mice generated *Cxcl12^+/KO^*mothers carrying wild-type, heterozygous, and knockout embryos. In parallel experiments, to identify the temporal requirements for CXCL12-CXCR4 signaling, we delivered a CXCR4 antagonist, AMD3100, to wild-type CD1 mice during specific developmental windows, blocking maternal CXCR4 activation while maintaining genetically normal trophoblasts. To understand how spiral artery growth and trophoblast invasion into the decidua might be affected by these perturbations, we used 2D tissue sections through the midline of the placenta, at E12.5, after spiral arteries are connected to the placenta and trophoblasts have enveloped arteries (Fig 1e, j, Extended Data Fig 1b).

Implantation sites with knockout embryos exhibited fewer decidual spiral arteries and invasive trophoblasts when compared to heterozygous controls (Fig 3b). Artery and trophoblast localizations were quantified in tissue sections by measuring their relative distance from the placental border (0) to myometrium (1) (Extended Data 4a). The results revealed that, with *Cxcl12^+/+^* and *Cxcl12^+/KO^* embryos artery-trophoblast spatial overlap was approximately 60%, whereas it was absent with knockout embryos (Fig 3c, d, Extended Data Fig 4b). This was independent of fetal viability, as all embryos were grossly normal, and we did not detect any lethality through genotyping ratios (Extended Data Fig 4c, d), consistent with the late gestational lethality of this line.^66,69^

For CXCR4 inhibition, AMD3100 was administered daily across four post-implantation periods: E4.5-11.5 (full), E4.5-6.5 (early), E6.5-8.5 (middle), and E8.5-11.5 (late) (Fig 3a). At E12.5, vehicle-treated deciduas were indistinguishable from genetic controls (e.g. *Cxcl12^+/+^* and *Cxcl12^+/KO^*), while full CXCR4 inhibition phenocopied *Cxcl12^KO/KO^* (Fig 3c, d and Extended Data Fig 4b). The shorter AMD3100 time windows revealed that artery and trophoblast behavior was particularly sensitive to inhibition during the early period. The early window histologically and quantitatively resembled *Cxcl12^KO/KO^* deciduas, reducing artery-trophoblast overlap to 1-6%, while the late period was indistinguishable from controls (Fig 3c, d, e, Extended Data Fig 4b). Aligning with the temporal gradient in spiral artery development (Fig 2b), the middle inhibition period demonstrated an intermediate phenotype, reducing artery-trophoblast overlap by approximately 50% (Fig 3c, d, Extended Data Fig 4b). Consistent with the requirement for CXCR4 signaling in development of fetal placental arteries,^51^ AMD3100 treatment from all treatment windows reduced placental thickness (Extended Data Fig 4e), yet none impacted embryo size (Extended Data Fig 4f). Maternal immune cells can utilize CXCR4 signaling (Extended Data 3b) and have been reported to require CXCR4 for successful pregnancy progression,^70,71^ yet CD45+ staining from genetic and pharmacological perturbations revealed no change in immune cell density in the decidua or placenta (Extended Data Fig 4g, h). This suggests that alterations in immune cell recruitment does not underlie the observed phenotypes.

Overall, these data establish that CXCL12-CXCR4 signaling during early pregnancy stages (E4.5-8.5) is critical for trophoblast invasion and proper artery development in the decidua throughout at least mid-gestation (E12.5), a defect that is independent of embryo growth.

### Loss of fetal Cxcl12 misdirects trophoblast migration and artery patterning

The near-complete absence of artery-trophoblast interactions observed in 2D sections appeared inconsistent with embryo survival, prompting a comprehensive 3D examination of the entire maternal-fetal interface. Whole-organ imaging of implantation sites from *Cxcl12^+/KO^* intercrosses confirmed that central optical sections recapitulated the 2D phenotype lacking arteries and trophoblasts (Fig 3f, left). However, upon examining the entire implantation site, it was apparent that the periphery was vastly different from the midline. In the periphery, the placenta of knockout embryos flanks a thinned decidua and thus is even closer to the myometrium than at the midline (Fig 3f, right). Atypically, invasive trophoblasts at the periphery were found within the myometrium and in the thin decidua, including enveloping a rarely-observed artery; in contrast, controls contained few trophoblasts and uterine arteries at periphery and none in the myometrium (Fig 3f, right). As before for wild type implantation sites (Fig 1l), we examined the 3D probability distribution of arteries and trophoblasts within the decidua and myometrium (see Methods). This confirmed that implantation sites containing knockout embryos had many peripherally localized trophoblasts, which resulted in decreased co-occurrence (Fig 3g).

Fully tracing spiral arteries in 3D images showed that they did connect to the placenta in knockouts, but at roughly one-quarter of the normal frequency (∼3 arteries in knockouts versus ∼12 in controls) (Fig 3h). Therefore, maternal blood flow into knockout placentas is likely reduced, but not to levels causing fetal demise (Extended Data Fig 4d). Quantifying trophoblast positioning across the entire decidua revealed a shift in cell localization: when compared to control, most trophoblasts were interstitial (70% in knockout vs 25% in control) as opposed to enveloping spiral arteries (10% in knockout vs 75% in control) (Fig 3i, j).

Most strikingly, the peripheral trophoblast invasion in knockouts extended into the myometrium, which was never seen in controls and is not normal for mouse pregnancy (Fig 3k). Approximately 20% of invading knockout trophoblasts migrated into the myometrium, and of these, ∼75% specifically targeted myometrial arteries (Fig 3j, k), demonstrating retained arterial tropism despite a redirection of invasion. Together, these 3D analyses reveal that loss of fetal *Cxcl12* causes mispatterning of the maternal-fetal interface: fewer arteries are present in the decidua and supply blood to the placenta, and a substantial fraction of trophoblasts over-invade into the myometrium, a phenotype reminiscent of placenta accreta.

### Disruption of CXCL12-CXCR4 models placental accreta

Because trophoblasts at unusually deep locations in the uterus occurs in placenta accreta in humans, we next asked whether CXCR4 signaling disruptions culminate in other hallmarks of this disease. Placenta accreta is predicted in patients during pregnancy when ultrasound detects direct attachment of the placenta to a thinned myometrium with a thin or absent intervening decidua and excessively large placental lacunae (maternal blood spaces).^19, 72^ A definitive diagnosis occurs after birth when parts of the placenta are not expelled and hemorrhage occurs. Direction adhesion of the uterus to itself or other external organs is sometimes observed in cases when surgery is performed. Histology of the adherent placenta also demonstrates abnormally deep trophoblast invasion into the myometrium.^12,20–23^ We investigated these same features in mice lacking CXCL12-CXCR4 signaling using both genetic and pharmacological manipulations.

Tissue sections at E12.5 revealed an average 50% reduction in decidual width alongside myometrial thinning in genetic knockouts and mice that received full, early, or middle CXCR4 inhibition, but not when AMD3100 was administered after E8.5 (Fig 4a, b). These results are consistent with the critical early window identified above (Fig 2c, d). Enlarged placental lacunae were also evident at E12.5 (Fig 4c). Machine learning-based quantification of lacunae volume in 3D images showed that they were on average 50% larger in *Cxcl12* knockout placentas (Fig 4d), resembling a key feature of placenta accreta.

**Figure 4:**
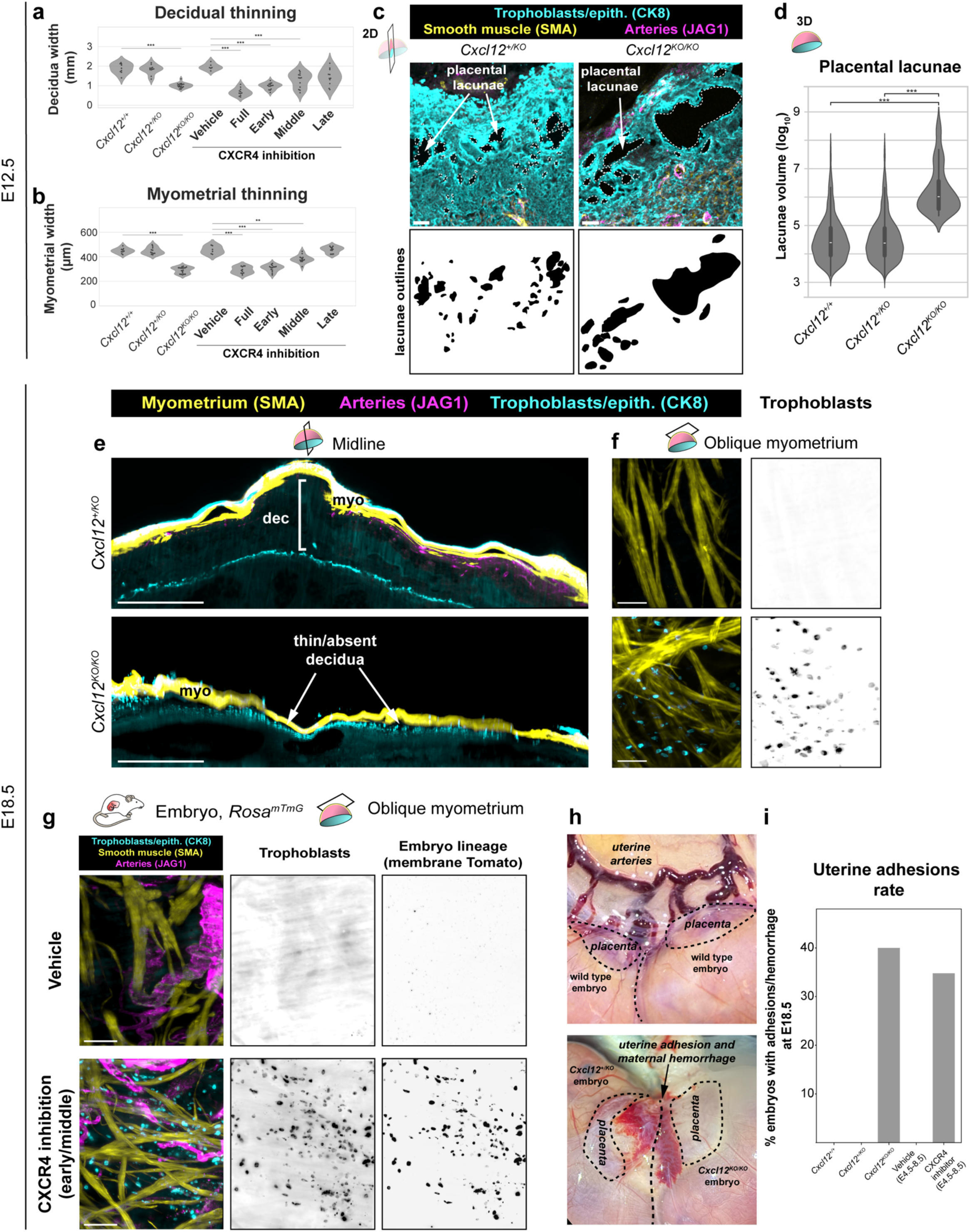
Disruption of CXCL12-CXCR4 models placenta accreta. **a, b,** Quantification of decidual (**a**), and myometrial thinning (**b**). Mice were analyzed at E12.5, *Cxcl12*^+/+^ (n = 9 implantation sites, n = 3 mice), *Cxcl12^+/KO^* (n = 11 implantation sites, n = 3 mice), *Cxcl12^KO/KO^* (n = 18 implantation sites, n = 5 mice), Vehicle (n = 6 implantation sites, n = 3 mice), AMD E4.5-11.5 (n = 10 implantation sites, n = 3 mice), AMD E4.5-6.5 (n = 14 implantation sites, n = 3 mice) AMD E6.5-8.5 (n = 13 implantation sites, n = 3 mice), AMD E8.5-11.5 (n = 11 implantation sites, n = 3 mice). **c**, Immunofluorescence of maternal blood lacunae from *Cxcl12^+/KO^* and *Cxcl12^KO/KO^* implantation sites at E12.5 (top), and outlines of the lacunae spaces (bottom). **d**, Quantification of maternal blood lacunae volume from *Cxcl12^+/KO^*intercrosses. Mice analyzed in 3D at E12.5, *Cxcl12^+/+^* (n = 5 implantation sites, n = 3 mice, *Cxcl12^+/KO^* (n = 9 implantation sites, n = 3 mice), *Cxcl12^KO/KO^* (n = 7 implantation sites, n = 3 mice). **e**, Optical longitudinal 60µm sections of E18.5 *Cxcl12^+/KO^* and *Cxcl12^KO/KO^* implantation sites. **f**, Optical 60µm sections of the oblique edge of the myometrium in E18.5 *Cxcl12^+/KO^* and *Cxcl12^KO/KO^* implantation sites. **g**, Optical 60µm sections from the myometrium of E18.5 implantation sites with vehicle treatment or CXCR4 inhibition in early + middle windows. **h**, Gross anatomical images of uterine adhesions in *Cxcl12^+/KO^* intercrosses at E18.5. **i**, Rate of macroscopically observed uterine adhesions. Mice analyzed at E18.5: *Cxcl12^+/+^* (n = 6 implantation sites, n = 3 mice), *Cxcl12^+/KO^* (n = 14 implantation sites, n = 3 mice), *Cxcl12^KO/KO^* (n = 5 implantation sites, n = 3 mice), Vehicle (n = 22 implantation sites, n = 2 mice), CXCR4 inhibition (n = 23 implantation sites, n = 2 mice). *P* value < 0.05 by Fisher’s Exact test and Bonferroni-correction. \*\*\**P* <0.001. *P* values from Kruskall-Wallis test with Bonferroni Correction (**a, b**) or Pairwise Mann-Whitney U Test (**d**). Scale bars, 100µm, oblique myometrium and lacunae; 500µm others. myo, myometrium; dec, decidua

To investigate whether these accreta-like symptoms became more severe in late pregnancy, the time when placenta accreta becomes an obstetric emergency in human patients, we performed 3D imaging of *Cxcl12* knockout implantation sites one day before birth (E18.5). Regions of knockout placentas were directly attached to the myometrium with a thin or completely absent intervening decidua (Fig 4e). Oblique 3D views capturing the myometrial layer showed trophoblast invasion present only in knockouts (Fig 4f). To confirm these CK8+ cells within the myometrium were indeed fetal trophoblasts, we performed crosses where the whole embryo expresses membrane Tomato while the mother is unlabeled (*Rosa26^mTmG^* males crossed to wild-type females). CXCR4 was then inhibited during the early + middle window (E4.5-8.5) with AMD3100, the previously-defined sensitivity window (Fig 3c, d). CK8+ trophoblast cells in the myometrium also stained for membrane Tomato, confirming fetal origin (Fig 4g). Note that this trophoblast over-invasion persisted to late gestation even when CXCR4 inhibition was restricted to early pregnancy, demonstrating that this brief signaling window has long-lasting consequences. Finally, gross observations of knockout and early + middle CXCR4 inhibitor uteri dissected at E18.5 revealed hemorrhages and uterine adhesions to adjacent mesometrium at a rate of 40% and 35%, respectively (Fig 4h, i).

Together, these results demonstrate that early disruption of CXCL12-CXCR4 signaling generates a mouse model that shares many features of human placenta accreta.

### Early CXCL12-CXCR4 inhibition disrupts decidualization

Having established that early CXCR4 inhibition later results in placenta accreta, we next sought to determine the temporal sequence of phenotypes in the days following implantation to identify the initial developmental insult and downstream progression of events. We first examined decidualization because *Cxcr4^Tango^; pTRE-H2B-GFP* mice indicated that uterine epithelium and stromal cells are responding to CXCL12 at early stages (Fig 2g-i), and transformation of the uterine epithelium and stroma is one of the earliest events in pregnancy, following implantation. Decidualization can be detected by upregulation of many markers, including Cadherin-3 (*Cdh3*) and Prolactin (*Prl8a2*).^4–8^ We immunostained E6.5 tissue sections of implantation sites from either *Cxcl12* knockout mice or early-window CXCR4 inhibition for CDH3. This revealed reduced CDH3 expression throughout the central uterine stroma with both perturbations, while some of the most peripheral anti-mesometrial stromal cells still expressed CDH3 (Fig 5a). Quantitative PCR of microdissected deciduas from the same conditions revealed approximately 50% decrease in *Prl8a2* expression with either fetal *Cxcl12* deletion or early CXCR4 inhibition (Fig 5b), indicating that decidualization impairment precedes the mid-gestation artery and trophoblast defects observed.

**Figure 5:**
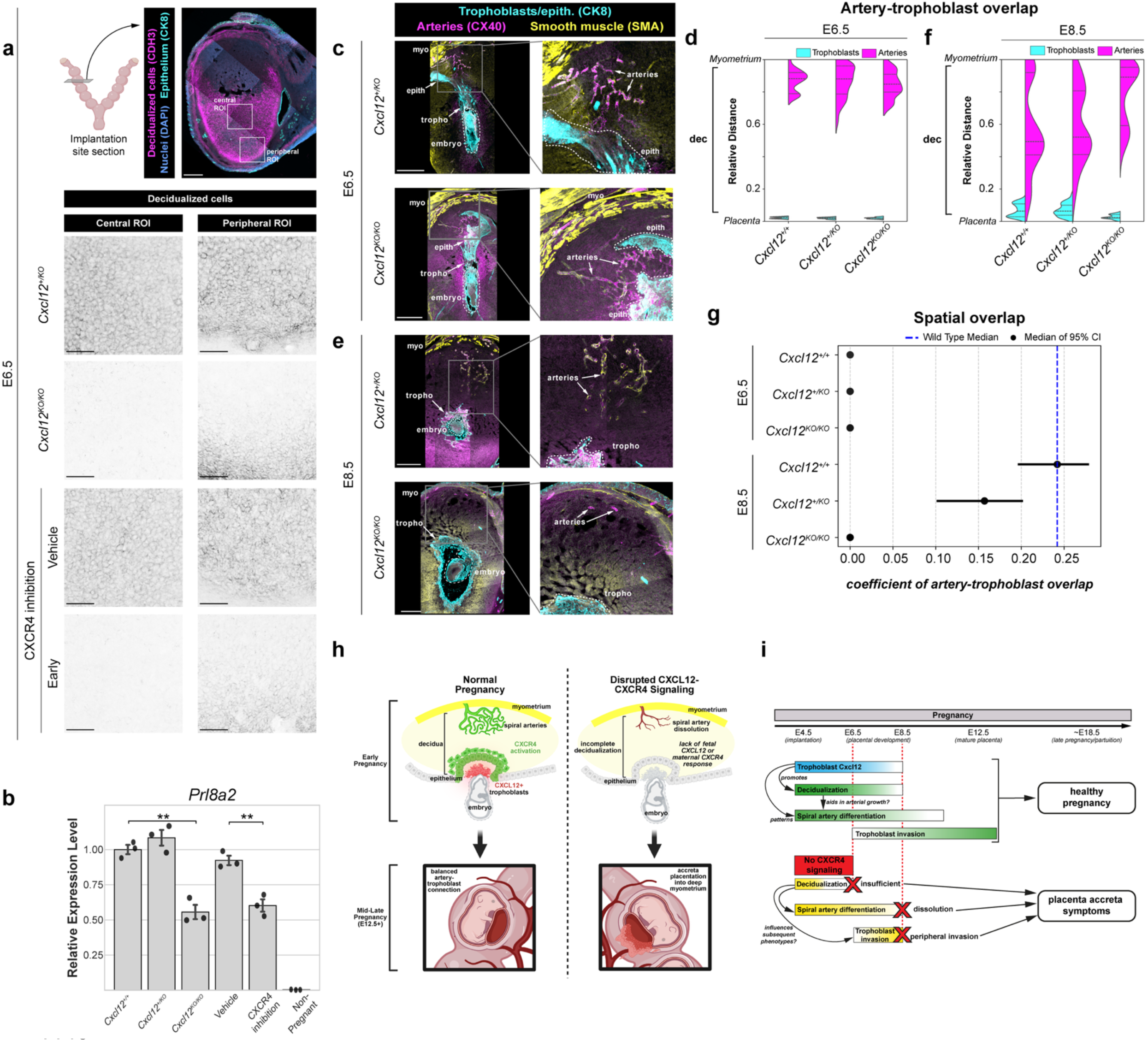
Disruption of CXCL12-CXCR4 causes a primary decidual defect. **a,** Representative anatomy and ROIs acquired at E6.5 for decidualization staining in genetic and pharmacological perturbations to CXCR4 signaling; central and peripheral ROIs from treatment groups. **b**, Real-time quantitative PCR measuring decidualization through normalized expression of *Prl8a2*. Gene expression was analyzed at E6.5 from implantation sites, *Cxcl12^+/+^* (n = 3 implantation sites, n = 2 mice, *Cxcl12^+/KO^* (n = 3 implantation sites, n = 2 mice), *Cxcl12^KO/KO^* (n = 3 implantation sites, n = 2 mice), Vehicle (n = 3 implantation sites, n = 2 mice), CXCR4 inhibition (n = 3 implantation sites, n = 2 mice); or in the non-pregnant uterus (n = 3 mice). Error bars, standard deviation. **c**, Immunofluorescence of *Cxcl12*^+/KO^ and *Cxcl12*^KO/KO^ E6.5 implantation sites. **d,** Relative positions of arteries and trophoblasts in the decidua normalized to the distance from the placenta to the uterine wall. Mice were analyzed at E6.5: *Cxcl12*^+/+^ (n = 8 implantation sites, n = 3 mice), *Cxcl12^+/KO^* (n = 7 implantation sites, n = 3 mice), *Cxcl12^KO/KO^* (n = 7 implantation sites, n = 5 mice). **e**, Immunofluorescence of *Cxcl12*^+/KO^ and *Cxcl12*^KO/KO^ E8.5 implantation sites showing disrupted development at E8.5 in *Cxcl12*^KO/KO^ implantation sites. **f**, Relative positions of arteries and trophoblasts in the decidua normalized to the distance from the placenta to the uterine wall. Mice were analyzed at E8.5: *Cxcl12*^+/+^ (n = 9 implantation sites, n = 3 mice), *Cxcl12^+/KO^* (n = 8 implantation sites, n = 3 mice), *Cxcl12^KO/KO^* (n = 7 implantation sites, n = 5 mice). **g**, Odds ratios with 95% confidence intervals for the interaction of artery and trophoblast interactions in the uterine wall from E6.5 and E8.5 implantation sites. **h, i**, Schematics for the role of CXCL12-CXCR4 signaling in early pregnancy and its ripple effects in the manifestation of phenotypes in later pregnancy. \*\**P*<0.01. *P* values from One Way ANOVA with Tukey’s Post-Hoc Test. Scale bars, 100µm, decidualization ROIs; 500µm, all others. myo, myometrium; epith, epithelium; dec, decidua; tropho, trophoblasts.

To determine when decidual artery defects first become apparent, we generated litters from *Cxcl12^+/KO^* intercrosses at E6.5 and E8.5 and quantified artery and trophoblast localization in 2D tissue sections of implantation sites. At E6.5 when we first detected an artery remodeling zone near the ectoplacental cone with little to no trophoblast invasion (Fig 1c, g), *Cxcl12* knockouts were indistinguishable from controls, indicating that artery development is initialized normally at this stage (Fig 5c, d, g).

However, at E8.5, spiral arteries near knockout placentas were diminished (Fig 5e, f, g). Thus, disruption of CXCL12-CXCR4 signaling triggers a temporal cascade: impaired decidualization as early as E6.5 precedes a block and/or reversal in artery development and decreased trophoblast invasion by E8.5, which we find ultimately culminates in placenta accreta at term, even when CXCR4 inhibition is released by E8.5.

## Discussion

Here, we utilize whole-organ imaging of the maternal-fetal interface in mice to investigate the dynamics of uterine spiral artery development and trophoblast invasion in 3D across gestation. Our results demonstrate that, by quantification of trophoblast and artery positioning in the whole implantation site, it can be appreciated that the mouse placenta is more invasive than previously thought. Thousands of trophoblasts invade hundreds of microns to fill the entire decidual thickness and extensively envelope the majority of maternal spiral arteries. This holistic quantification revealed why previous studies underestimated mouse invasion. Trophoblasts consistently envelope 10-12 spiral arteries by mid-gestation, but this overlap can occur at different regions within the 3D space of the implantation site such that 2D sections at the direct midline don’t always regularly capture the extent of invasion. Armed with this imaging technique, we were able to assess the specific developmental defects occurring in the absence of CXCL12-CXCR4 signaling and detect many hallmarks used to predict and diagnose human placenta accreta.

Using 3D imaging, along with the tissue sectioning methods standard to the field, we established that proper decidualization and persistence of spiral arteries during early uteroplacental development requires fetal *Cxcl12* and CXCR4 signaling (Fig 5h, i).

Trophoblast *Cxcl12* expression and activation of CXCR4 in decidual stroma and arteries overlaps with decidualization and artery development time points as shown by our lineage tracing. Both genetic deletion of fetal *Cxcl12* and early pharmacological CXCR4 inhibition results in reduced expression of decidualization markers at E6.5 and fewer spiral arteries at E8.5. Subsequently, at mid-gestation, implantation sites contain roughly one-quarter of the expected number of arteries connected to the placenta, coupled with aberrant trophoblast invasion of the myometrium. At parturition, these massive structural defects cause a number of placenta accreta-like features seen in humans, including a thin or absent decidua, thin myometrium, expansive placental lacunae, and trophoblast invasion into the myometrium with hemorrhages and adhesions to external organs. Thus, our data defines an early window post-implantation in which a molecular pathway can be disrupted to result in continued embryonic growth but, pathological placentation to the detriment of the mother.

Placenta accreta, is heavily associated with uterine fibrosis and altered proteoglycan expression–hallmarks of prior cesarean scarring.^73–77^ Thus, it is reasonable to hypothesize that the alterations to CXCL12 signaling gradients by disrupted extracellular matrix (ECM) could result in pathological trophoblast invasion. Because CXCL12 exhibits a strong biochemical affinity for proteoglycans,^78–80^ aberrant ECM composition, such as that in cesarean scars, could physically trap CXCL12 within fibrotic tissue. This sequestration would disrupt the highly directional CXCR4 signaling required for normal decidualization and spiral artery recruitment, offering a parsimonious mechanism for placenta accreta pathogenesis, in both scarred and unscarred contexts.

The pathogenesis in our system is temporally preceded by early decidual defects, the insult thought to contribute to the human disease. Proper decidualization requires a tightly coordinated network of maternal and fetal signals;^4,81–83^ *in vivo*, and our data suggests that fetoplacental CXCL12 signaling to maternal epithelial and stroma influences uterine gene expression, potentially through the induction of the known required factor *Lif*, to promote uteroplacental development.^84–87^ Interestingly, *Cxcl12* may be a target of the broad epigenetic reprogramming crucial for decidualization, including the EZH2-dependent H3K27me3 silencing in decidual stromal cells.^88–90^ Maternal epigenetic silencing of chemokine genes including *Cxcl12* aligns with our observation that early pregnancy relies specifically on fetoplacental *Cxcl12*, suggesting that spatiotemporal restriction of the CXCL12-CXCR4 axis may ensure proper demarcation of the decidual boundary against placental overgrowth.

Our lineage tracing provides insight into how spiral arteries form; we show that spiral arteries directly differentiate from capillary ECs. We propose these newly differentiating artery ECs coalesce to form arterial vessels *in situ* that connect with pre-existing myometrial arteries, similar to how coronary arteries differentiate and connect with the aorta, guided by blood flow.^39^ We see early spiral artery development initiating at E6.5 in *Cxcl12* knockouts but with later dissolution, resulting in a stunted maternal arterial supply and a 4-fold decrease in arteries connecting to the placenta by mid-gestation. We show that CXCL12-CXCR4 is critical during the E4.5-8.5 period; however, our lineage tracing indicates artery differentiation still occurs outside of this window, suggesting compensatory signals may exist. Potential pathways previously implicated in vascular growth during periods surrounding E4.5-8.5 include P4-PR and VEGFA-VEGFR2 signaling, which is known to drive vascular growth and permeability in the pre-and peri-implantation period^91–94^ and VEGFC-VEGFR3, which is required for arteries after mid-gestation with knockout causing defects beginning at E13.5.^95^ Thus, spiral arteries could be regulated by a temporal hand-off between signaling axes throughout pregnancy.

The dramatic maternal-fetal interface phenotype caused by altered CXCL12-highlights its potential importance. It is tempting with current transcriptomic analyses of human pregnancy to speculate that predicted CXCL12-CXCR4-mediated interactions between trophoblasts and endothelial cells are functioning similarly to what we observe here.^52,53^ While sequencing of placenta accreta reveals alterations in related networks – such as regulation of angiogenesis, CXCL signaling, and hypoxia responses^96,97^ – human genome-wide association studies have not yet been performed for placenta accreta. Previous human genomic association studies have not linked *CXCL12* to pregnancy complications such as preeclampsia and gestational hypertension, but these studies are among the first reports and may require additional power to finely map deleterious genetic loci.^98–101^ Given that *CXCL12* variants are common in the population, future studies may identify associations with pregnancy complications, as was recently discovered for coronary artery disease and patterning.^46^

The establishment of this deeply invasive murine model opens new avenues for placenta accreta modeling and broader understanding of uteroplacental development. Having redefined this structure using 3D imaging, we can now leverage this anatomical framework to investigate trophoblast invasion, vascular remodeling, and spatial dynamics utilizing the robust genetic toolkit present in mice. A critical next step is interrogating CXCR4 signaling in early pregnancy to dissect early molecular drivers of uteroplacental development. Combining our 3D imaging with targeted molecular profiling, future studies can elucidate previously undiscovered pathophysiology underlying placenta accreta and identify potential early biomarkers of pathological trophoblast invasion. In conclusion, our whole-organ imaging approach provides a novel framework for murine placental development, redefining the spatial extent of trophoblast invasion and maternal spiral artery envelopment. By identifying the critical fetal-maternal CXCL12-CXCR4 signaling axis, this work elucidates the basic molecular mechanisms governing invasive placentation, spiral artery differentiation, and the structural integrity of the maternal-fetal interface.

## Methods

### Mouse models

All mouse husbandry and experiments were performed in accordance with Stanford Institutional Animal Care and Use Committee (IACUC) guidelines. Animals were housed in a pathogen-free facility, with a maximum of 5 adult mice per cage. Animals were monitored daily and kept under a 12-hour light/dark cycle. All mice in this study were maintained on a mixed background, and have been previously described: CD1 (Charles River Laboratories, Strain Code #022), *ROSA26^tdTomato^* (Jackson Laboratory, Stock #007909), *ROSA26^mTmG^* (Jackson Laboratory, Stock #007576), *Cxcl12^dsRed^* (Jackson Laboratory, Stock #022458), *pTRE-H2B-GFP* (Jackson Laboratory, Stock #005104), *Cx40-CreER*,^102^ *Cxcr4^Tango^,*^64^ *AplnrCreER*.^103^

### Breeding, tamoxifen, doxycycline, and compound administration

Timed pregnancies were determined through the presence of a vaginal plug on the morning day of, which was designated as embryonic day E0.5. To activate inducible *CreER* lines, tamoxifen (Sigma Aldrich, T5648-5G) in corn oil (Sigma Aldrich, C8267-500mL) was injected intraperitoneally at 75mg/kg. For pharmacological inhibition of CXCR4, AMD3100 (Selleck Chemicals, S8030) was dissolved in PBS and injected at 40mg/kg at noted days of pregnancy. Injection with PBS was used as a vehicle control.

### Immunofluorescence

All tissues were dissected in ice cold PBS (Cytiva, SH3025601) and fixed in 4% PFA (Electron Microscopy Sciences, 15714) for 1 hour, rotating at 4°C. Following fixation, tissues were washed 3x15 minutes in PBS, then stored in PBS with 0.01% sodium azide (w/v, Sigma Aldrich, 71289-5G) until processed. Tissues were placed in 30% sucrose (Sigma Aldrich, S0389-1KG) at 4°C overnight. The following day, tissues were embedded in OCT (Thermo Fisher, 23-730-571), snap frozen, and stored at -20°C. Cryosections were made at 70µm thickness onto microscope slides (Thermo Fisher, 12-550-15) and stored at -20°C until further processing. Slides were thawed at room temperature for 15 minutes in a humidified incubation chamber, followed by 3x15 minute washes in PBS-T (PBS with 0.05% Tween-20 (Sigma Aldrich, P1379-500mL)), followed by incubation in a blocking solution of 5% normal donkey serum (Jackson ImmunoResearch, 102644-006) prepared in PBS-T for 1 hour at room temperature.

Primary antibodies were diluted in blocking solution and incubated overnight at 4°C. Primary antibodies used were: rat anti-CK8 (DSHB, TROMA-I, 1:100), mouse anti SMA-Cy3 (Sigma, C6198-.2mL, 1:500), goat anti-JAG1 (R&D, AF599, 1:500), rabbit anti-CX40 (Alpha Diagnostics, CX40-A, 1:250), rabbit anti-ERG (Abcam, ab92513, 1:300), goat anti-CDH3 (R&D, AF761, 1:250). The following day, slides were washed in PBS-T for 3x30 minutes and incubated with secondary antibodies and DAPI (Sigma Aldrich, D9542-1MG) nuclear counterstain diluted in blocking solution at room temperature for one hour. Secondary antibodies used were: Alexa Fluor donkey anti-rat 488 (Invitrogen, A48269, 1:250), Alexa Fluor donkey anti-goat 488 (Invitrogen, A32814, 1:250), Alexa Fluor donkey anti-rabbit 594 (Invitrogen, A32754, 1:250), Alexa Fluor donkey anti-rat 594 (Invitrogen, A48271, 1:250), Alexa Fluor donkey anti-rabbit 647 (Invitrogen, A32795, 1:250), Alexa Fluor donkey anti-goat 647 (Invitrogen, A32849, 1:250), Alexa Fluor donkey anti-rat 647 (Invitrogen, A48272, 1:250). Slides were washed in PBS-T for 3x30 minutes, then mounted in Fluoromount G (SouthernBiotech, 0100-01) and sealed with nail polish prior to confocal imaging. Imaging was performed using a Zeiss LSM-980 confocal microscope (10x or 20x objective lens) with ZEN 3 software (Zeiss).

### Whole-Organ Clearing and Immunolabeling

All staining was performed using a modified DISCO-based protocol^104^ for light sheet imaging. Briefly, all tissues were dissected in ice cold PBS and fixed in 4% PFA for 1 hour, rotating at 4°C. Following fixation, tissues were washed 3x15 minutes in PBS, then stored in PBS with 0.01% sodium azide (w/v) until ready for processing. Tissues were dehydrated in a methanol (Sigma Aldrich, 179337-6x1L) series (20%/40%/60%/80%/100%/100%), prepared in B1N solution (0.3M glycine (Thermo Fisher, 15527013) 0.1% Triton X-100 (v/v, Sigma Aldrich, T8787-250mL) 0.01% sodium azide (w/v), 0.001N NaOH (Thermo Fisher, BP359-500)) each for 1 hour at room temperature, with gentle rocking. Following dehydration, samples were rocked overnight in 100% dichloromethane (Thermo Fisher, AA47352M1) at room temperature, under a fume hood. The next day, tissues were washed for 4 hours in 100% methanol at room temperature rocking. Tissues were then placed into 10% hydrogen peroxide (Sigma Aldrich, 216753-500mL) prepared in methanol, shaking at 4°C, inside of a photobleaching chamber; tissues were photobleached until tissues were pale white, approximately 3 days. Tissues were rehydrated in a methanol series (80%/60%/40%/20%) prepared in B1N solution, for 1 hour each, rocking at room temperature. They were then washed for 1 hour in B1N solution, followed by fresh B1N solution overnight rocking at room temperature. The following morning, tissues were washed 1 x 2 hours in PTwH (PBS with 0.2% Tween-20, 0.1% heparin (Sigma Aldrich, H3393-10KU), 0.01% sodium azide (w/v)), then incubated with primary antibodies prepared in PTwH with 5% DMSO, and 3% NDS. Tissues were incubated with primary antibodies and gentle rocking at 4°C for two days, followed by two days at room temperature, and up to two weeks at 37°C, dependent upon sample size. Primary antibodies used were: rabbit anti-RFP (Rockland, 600-401-379, 1:500), rat anti-CK8 (DSHB, TROMA-I, 1:100), mouse anti SMA-Cy3 (Sigma Aldrich, C6198-.2mL, 1:750), goat anti-JAG1 (R&D, AF599, 1:500). Following incubation, samples were washed 1 x 30 minutes, then 5 x 1.5 hours in PTwH at room temperature rocking; for whole uterus samples, these washes were extended in frequency and time over two days. Tissues were then incubated with secondary antibodies prepared in PTwH with 3% NDS at the same temperatures and timing as primary antibodies. Secondary antibodies used were: Alexa Fluor donkey anti-rat 488 (Invitrogen, A48269, 1:250), Alexa Fluor donkey anti-goat 647 (Invitrogen, A32849, 1:250), Alexa Fluor donkey anti-rabbit 647 (Invitrogen, A32795, 1:250), Alexa Fluor 790 donkey anti-goat (Jackson Immuno, 705-655-147, 1:250). Following secondary antibody incubation, tissues were washed 1 x 30 minutes and 5 x 1.5 hours with PTwH, followed by fresh PTwH overnight at room temperature; for whole uterus samples, these washes were extended in frequency and time over two days. Tissues were embedded in blocks of 2% agarose (Thermo Fisher, BP160-500) prepared in PBS. Sample blocks were dehydrated in a methanol series (20%/40%/60%/80%/100%/100%) prepared in PBS, for 1 hour each at room temperature rocking. The last 100% methanol wash was left overnight, and the next day was replaced with 100% dichloromethane under a fume hood for the entire day.

Samples were then placed in new tubes with ethyl cinnamate (Sigma Aldrich, 112372-100G) inverted 3-4 times, then stored overnight for tissue clearing. All samples were imaged using a Miltenyi UltraMicroscope Blaze (4x or 12x objective) with ImSpector 7.8.0 Software (Miltenyi).

### RNA extraction and quantitative PCR

Tissue samples for RNA extraction were dissected in ice cold PBS and snap frozen. Tissue was initially homogenized using a cooled ceramic mortar and pestle (Thermo Fisher, FB-961B & FB-961-K), and were placed in ZR BashingBead Lysis Tubes (Zymo, S6003-50) containing TRIzol (Thermo Fisher, 15596026) and 80U/ml Proteinase K (New England Biolabs, P8107S). Tissue was further homogenized using a Qiagen TissueLyser on a 5x30s protocol with 5 seconds rest between runs, then placed on ice for 5 minutes. Samples were then incubated at 37°C and 1000RPM for 30 minutes, then spun at 12,000 g for 5 minutes at 4°C. 750µl of supernatant was transferred to a new tube with 750µl ethanol (Fisher BP2818500). Samples were cleaned up using RNA Clean & Concentrator (Zymo, R1015), followed by cDNA synthesis using Verso cDNA Synthesis Kit (Thermo Fisher, AB1453B). Quantitative PCR with reverse transcription was performed on a QuantStudio 7 Flex Real-Time PCR system using PowerUP SYBR Green Master Mix (Thermo Fisher, A25778), and primers: *Actb*-F (GGCTGTATTCCCCTCCATCG), *Actb*-R (CCAGTTGGTAACAATGCCATGT), *Prl8a2*-F (GAGTCAACCTCACTTCTGGGC), and *Prl8a2*-R (CTGAGCAGCCATTCTCTCCT).

Genes were measured using technical triplicates taken from at least three independent uteri/implantation sites as biological replicates. Gene expression was calculated using the ΔΔCt method, normalized to a housekeeping gene, and relative sample expression normalization to *Cxcl12^+/+^* samples.

### Image processing and quantifications

Quantification of confocal images was performed using FIJI (NIH). A minimum of three implantation site sections within 240µm of the canal region of the placenta (E12.5 implantation sites) was used for quantification, at least two sections containing the ectoplacental cone (E6.5 implantation sites), or a minimum of three implantation site sections within 180µm of the ectoplacental cone tip (E8.5 implantation sites), per implantation site analyzed. Artery cells were quantified as ERG+ as well as JAG1+ or CX40+. Artery cells containing reporters (*Cxcr4^Tango^* or Cre lineage trace reporter) were quantified as ERG+, JAG1+ or CX40+, and reporter positive (GFP+ or tdTomato+).

Normalized distance to the uterine wall was normalized by defining the maximum Euclidean distance from the CK8+ placenta to the SMA+ myometrium as a value of 1; minimum Euclidean distance of CK8+ trophoblasts and CX40+ or JAG1+ artery lumens from the CK8+ placental border was used for quantification.

Quantification of light sheet images was performed using Imaris 10.2 (Oxford Instruments). The Imaris machine learning segmentation was used to create a surface of the SMA channel to represent the myometrium of the uterus; segmentation of the JAG1 channel represented arteries, CK8 segmentation represented trophoblasts.

Machine learning segmentation of maternal blood spaces was performed by training on lacunae regions lacking CK8 reactivity within the placenta. Quantification of the localization of trophoblasts was performed using Imaris’s slice function, where CK8+ puncta were manually counted in every z-section from every implantation from all pregnancies analyzed. CK8+ trophoblasts within/surrounding JAG1+ arteries were defined as spiral artery-adjacent; interstitial trophoblasts were defined as CK8+ puncta within the decidua, but not within a JAG1+ artery. CK8+ puncta within the SMA+ myometrium were quantified as within the myometrial stroma (not within a JAG1+ artery), myometrial artery-adjacent (within the SMA+ myometrium, and within a JAG1+). To generate 3D probability densities, Imaris’s spots function was used on the CK8 and JAG1 channel to generate locations of signal for arteries and trophoblasts. Volume-based filtering of erroneous spots below the size detection of the light sheet microscope was applied. All spots were removed from analysis within the region of mesometrial CK8 reactivity surrounding the myometrium. The position of each spot was exported from Imaris, and for each implantation site, the X,Y,Z positions of anatomical referents (luminal epithelium of the inter-implantation sites, outer edge of the myometrium, outer edge of the junctional zone, and “front” and “back” of the implantation site) were extracted to normalize values on a (0,1) axis within a half-dome region.

### Statistics and reproducibility

No statistical method was used to predetermine sample size. All experiments were carried out with at least one biological replicate litter. Number of animals used, and statistical significance are described in the corresponding figure legends.

*P* values were generated from One Way ANOVA with Tukey Post-Hoc Test, Independent t-test, Pairwise Mann-Whitney U Test, or Kolmogorov-Smirnov Test with Dunn Post-Hoc Test. Multiple hypothesis testing was performed using Bonferroni corrections where applicable. Statistical analyses are denoted in figure legends. Spatial distributions of arteries and trophoblasts were modeled using Kernel Density Estimates. To quantify the spatial overlap between these populations, the Bhattacharyya coefficient (BC) was calculated by numerically integrating the square root of the product of their respective density functions. Statistical uncertainty was assessed via bootstrapping with replacement (n = 1000) to determine the median BC and 95% confidence intervals. Contingency table data was analyzed using a Fisher’s exact test followed by a post-hoc Bonferroni correction.

### Reporting Summary

Further information on research design is available in the Nature Portfolio Reporting Summary linked to this article.

## Data Availability

Any additional information required to reanalyze the data reported or replicate experiments performed in this paper is available from the lead contact upon request.

## Code Availability

All computational analyses were performed using Python 3.12.2 and are available on GitHub at https://github.com/james-b-z/CXCL12-placenta.

## Acknowledgements

Grant support: K.R.-H, NIH/NHLBI (R01-HL128503 and R01-HL171326), Stanford Dunlevie Maternal-Fetal Medicine Center Seed Grant, they are an HHMI investigator; J.B.Z, NIGMS, NIH (T32GM007276) and NSF-GRFP (DGE-1656518); M.N.M. is a Stanford Science Fellow and is supported with K.R.-H in part by grant number GV673607342 from Coefficient Giving. We thank Susan Fisher for helpful experimental suggestions, and Susan Fisher, Dominique Bergmann, Virginia Winn, Jeffrey Naftaly, and Azalia Martínez Jaimes for manuscript comments.

## Contributions

All authors were involved in study conceptualization and design, data acquisition, analysis, and interpretation of data, drafting and critical revision of the manuscript.

## Ethics declarations

### Competing interests

The authors declare no competing interests.

**Supplementary Video 1 – E6.5 implantation site**

A representative video showing 3D reconstructions of an E6.5 implantation site imaged using light-sheet microscopy. CK8 (cyan) indicates trophoblasts and epithelium, JAG1 (magenta) indicates arteries, and SMA (yellow) indicates smooth muscle, optical section is 50µm in width.

**Supplementary Video 2 – E8.5 implantation site**

A representative video showing 3D reconstructions of an E8.5 implantation site imaged using light-sheet microscopy. CK8 (cyan) indicates trophoblasts and epithelium, JAG1 (magenta) indicates arteries, and SMA (yellow) indicates smooth muscle, optical section is 50µm in width.

**Supplementary Video 3 – E12.5 implantation site**

A representative video showing 3D reconstructions of an E12.5 implantation site imaged using light-sheet microscopy. CK8 (cyan) indicates trophoblasts and epithelium, JAG1 (magenta) indicates arteries, and SMA (yellow) indicates smooth muscle, optical section is 50µm in width.

**Supplementary Video 4 – 3D probability distribution overview video**

A representative video rotating 3D reconstruction of an E12.5 implantation site imaged using light-sheet microscopy. CK8 (cyan) indicates trophoblasts and epithelium, JAG1 (magenta) indicates arteries, and SMA (yellow) indicates smooth muscle. Rotation to show half-dome-like shape of implantation site, followed by optical section slices of 50µm in width through the XZ plane. Red and green dots placed on optical section to show potential spots analysis for a single plane, whose positions are extracted for analysis.

**Supplementary Video 5 – E12.5 *Cxcl12^KO/KO^* implantation site**

A representative video showing 3D reconstructions of an E12.5 *Cxcl12^KO/KO^*implantation site imaged using light-sheet microscopy. CK8 (cyan) indicates trophoblasts and epithelium, JAG1 (magenta) indicates arteries, and SMA (yellow) indicates smooth muscle, optical section is 50µm in width.

---

**Extended Data Fig. 1:**
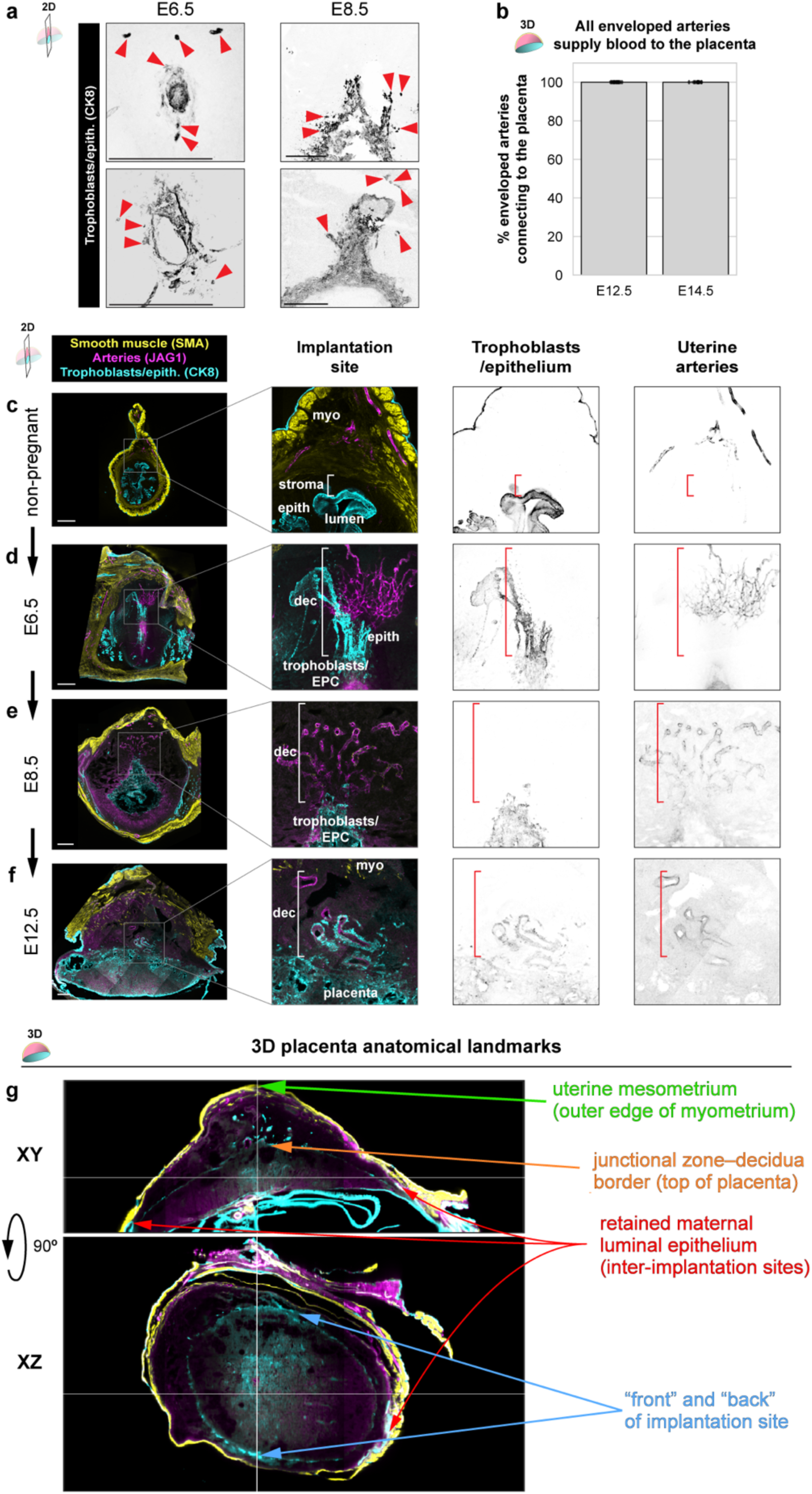
Two-dimensional imaging and quantification of placental development. **a,** Immunofluorescence of E6.5 (left) and E8.5 (right) implantation sites showing sparse trophoblast invasion (arrowheads). **b**, Quantification of the number of enveloped arteries connecting to the placenta through whole-organ imaging and light-sheet microscopy. Mice were analyzed at E12.5 (n = 13 implantation sites, n = 4 mice), and E14.5 (n = 12 implantation sites, n = 3 mice). **c-f**, Immunofluorescence of non-pregnant uterus (**c**), E6.5 implantation site (**d**), E8.5 implantation site (**e**), and E12.5 placenta (**f**) cross-sections to visualize arteries and trophoblasts. **g**, XY and XZ cross-sectional views of an E12.5 implantation site showing anatomical landmarks used for 3D density mapping. Scale bars, 500µm.

**Extended Data Fig. 2:**
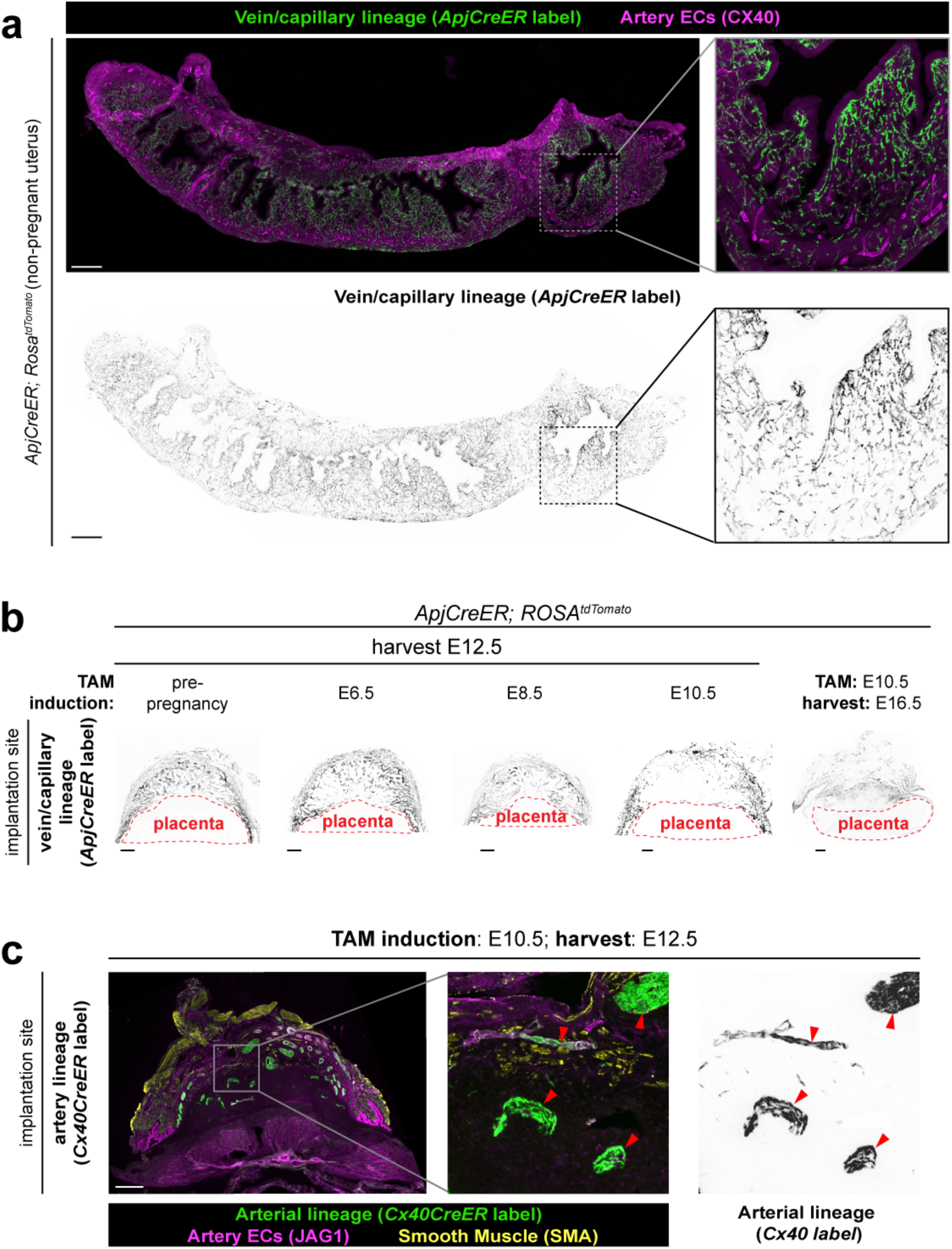
Lineage tracing of spiral arteries across gestation. **a**, Immunofluorescence of *ApjCreER* lineage labeled cells in the non-pregnant uterus one day after tamoxifen induction. **b**, Immunofluorescence of *ApjCreER* lineage labeled cells in the whole implantation site at noted days of induction and harvest. **c**, Immunofluorescence of *Cx40CreER* lineage labeled cells in the E12.5 implantation site after E10.5 induction. Arrowheads denote traced spiral arteries with retained lineage labeling. Scale bars, 500µm.

**Extended Data Fig. 3:**
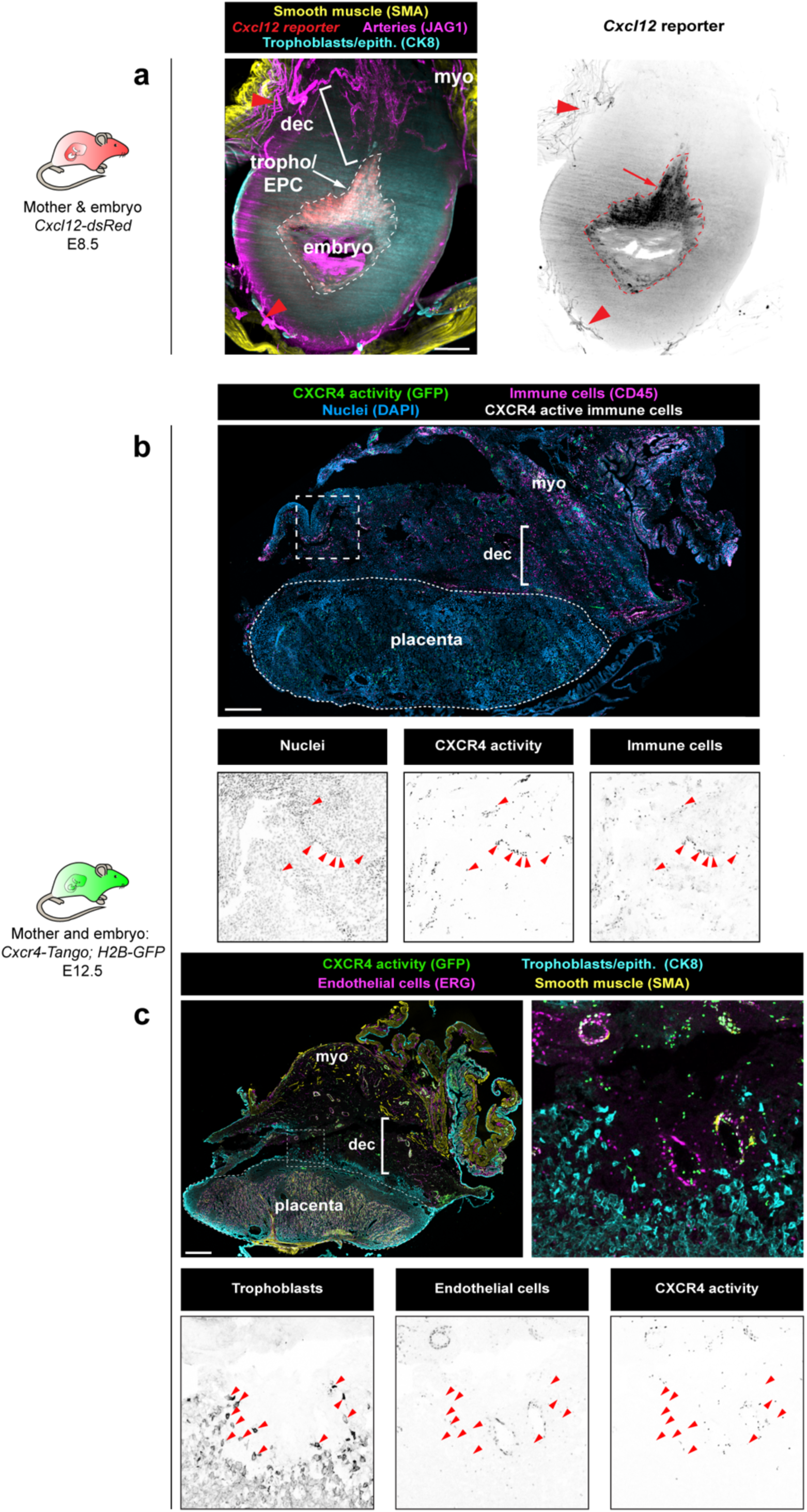
CXCL12-CXCR4 during uteroplacental development. **a**, Representative cross-sectional 60µm optical section from whole-organ light sheet imaging of *Cxcl12^dsRed^* at E8.5. Arrow, trophoblastic *Cxcl12* expression; arrowheads, myometrial artery *Cxcl12* expression. **b**, CXCR4 activation in the E12.5 implantation site is present on immune cells in the decidua. **c**, CXCR4 activation in the E12.5 implantation site is not activated on trophoblasts. Scale bars, 500µm. myo, myometrium; dec, decidua; tropho/EPC; trophoblasts/ectoplacental cone

**Extended Data Fig. 4:**
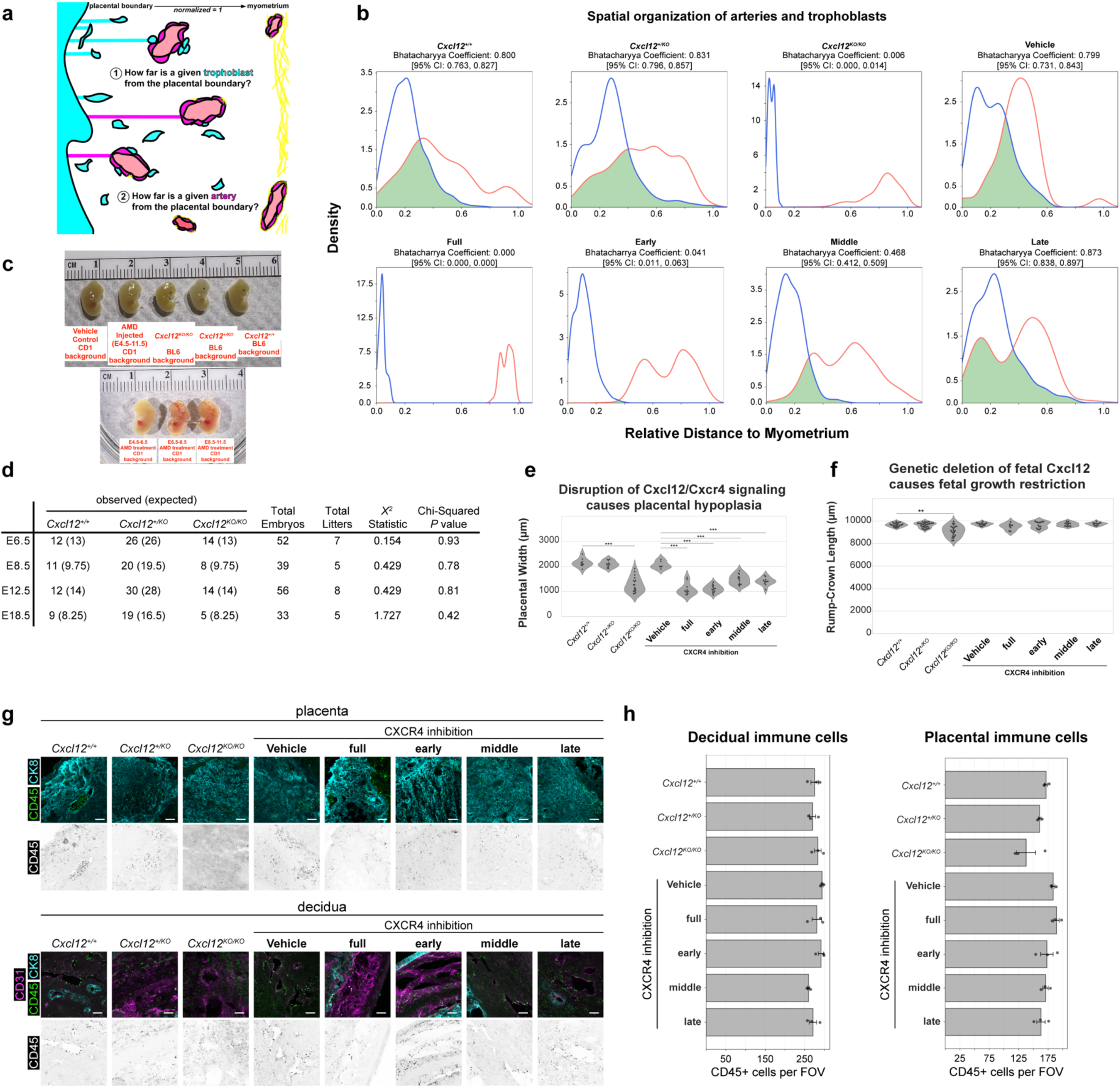
Further 2D phenotyping of CXCL12-CXCR4 perturbations. **a**, Schematic for 2D quantification of relative position of arteries and trophoblasts within the uterine wall. **b**, Kernel density estimates graphs for artery and trophoblast interaction at E12.5. **c**, Gross anatomical pictures of embryos harvested at E12.5 from denoted genetic or pharmacological perturbations. **d**, Chi-squared analysis on genotyping of embryos harvested from *Cxcl12^+/KO^* intercrosses. **e**, **f**, Quantification of placental hypoplasia (**e**), fetal growth restriction (**f**). Mice were analyzed at E12.5, *Cxcl12*^+/+^ (n = 9 implantation sites, n = 3 mice), *Cxcl12^+/KO^* (n = 11 implantation sites, n = 3 mice), *Cxcl12^KO/KO^* (n = 18 implantation sites, n = 5 mice), Vehicle (n = 6 implantation sites, n = 3 mice), AMD E4.5-11.5 (n = 10 implantation sites, n = 3 mice), AMD E4.5-6.5 (n = 14 implantation sites, n = 3 mice) AMD E6.5-8.5 (n = 13 implantation sites, n = 3 mice), AMD E8.5-11.5 (n = 11 implantation sites, n = 3 mice). **g**, Immunofluorescence of immune cells within the placenta (top) and decidua (bottom) at E12.5. **h**, Quantification of immune cells per field-of-view from the placenta and decidua of denoted treatment groups. \*\**P* <0.01, \*\*\**P* <0.001. *P* values from Kruskal-Wallis test with Bonferroni Correction. Scale bars, 100µm.

